# Interplay of the Trypanosome Lytic Factor and innate immune cells in the resolution of cutaneous *Leishmania* infection

**DOI:** 10.1101/2020.07.02.184358

**Authors:** Jyoti Pant, Marie Samanovic, Maria T Nelson, Mert K Keceli, Joseph Verdi, Stephen M. Beverley, Jayne Raper

## Abstract

Trypanosome Lytic Factor (TLF) is a primate-specific high-density lipoprotein complex that contains APOL1, the lytic component. Human TLF confers sterile immunity to many animal-infective extracellular *Trypanosoma* Ssp, which have been extensively investigated. Here, we have dissected the underappreciated role of TLF and neutrophils against intracellular *Leishmania* in intradermal infection. Our data show that mice producing human or baboon TLF have reduced parasite burdens when infected intradermally with metacyclic promastigotes of *L. major*. This TLF-mediated reduction in parasite burden was lost in neutrophil-depleted TLF mice, suggesting that early recruitment of neutrophils is required for TLF-mediated killing of *L. major*. Neutrophils and macrophages are the predominant phagocytes recruited to the site of infection. Our data show that acidification of the macrophage phagosome is essential for TLF-mediated lysis of metacyclic promastigotes. *In vitro* we find that only metacyclic promastigotes co-incubated with TLF in an acidic milieu were lysed. However, amastigotes were not killed by TLF at any pH. These findings correlated with binding experiments, revealing that labeled TLF binds specifically to the surface of metacyclic promastigotes, but not to amastigotes. During differentiation to the amastigote stage, the parasites shed their surface glycoconjugates. Metacyclic promastigotes of *L. major* deficient in the synthesis of surface glycoconjugates (*lpg1*^*-*^ and *lpg5A*^*-*^*/lpg5B*^*-*^) were partially resistant to TLF lysis. We propose that TLF binds to the outer surface glycoconjugates of metacyclic promastigotes, whereupon APOL1 forms a pH-gated ion channel in the plasma membrane, resulting in osmotic lysis. We hypothesize that resistance to TLF requires shedding of the surface glycoconjugates, which occurs upon phagocytosis by immune cells.

**Author Summary:** Leishmaniasis is a common term used for disease caused by parasites of the genus *Leishmania.* Depending on the parasite species and the clinical outcome of the disease, leishmaniasis can be divided into cutaneous, muco-cutaneous and visceral. Of the three, cutaneous leishmaniasis is the most common form, which is usually characterized by a localized lesion due to the infection of immune cells, primarily macrophages of the dermis and local lymph nodes. Sometimes, infected individuals can remain asymptomatic and do not show visible lesions. Moreover, the time between the infection and appearance of lesions are also variable and range from a few weeks to months and a few years in some cases. This subclinical stage of leishmaniasis depends on a variety of factors: parasite virulence, infectious dose, and host immune response. Therefore, it is important to understand the host-parasite interaction and its role in the clinical outcome of the disease. Here, we analyze the interaction between a cutaneous strain of *Leishmania* and a host innate immune factor called Trypanosome Lytic Factor (TLF). TLF is a type of High-Density Lipoprotein (HDL) complex that circulates in our plasma. TLF kills extracellular African Trypanosomes by lysing the parasites. The lytic ability of TLF is due to the primate specific protein APOL1 that forms pH gated ion channels. APOL1 inserts into biological membranes at acidic pH and forms a closed ion-channel that opens when the membrane associated APOL1 is exposed to neutral pH.

Using transgenic mice producing primate TLF, we show both human and baboon TLFs ameliorate cutaneous *Leishmania major* infection. The reduction in parasite burden correlated with: 1. infectious dose of metacyclic promastigotes and 2. the concentration of circulating TLF in mouse plasma. The early recruitment of neutrophils at the site of infection was required for the reduction of parasite burden by TLF. Macrophages, another major cell that phagocytoses metacyclic promastigotes at the site of infection require an acidified phagosome for TLF mediated killing of *L. major.* The acidification step is also essential for TLF mediated lysis of axenic metacyclic promastigotes of *Leishmania in vitro.* The susceptibility of metacyclic promastigotes to TLF mediated lysis is governed by the surface glycoconjugates of *Leishmania*. We find that surface glycoconjugate deficient *Leishmania* are resistant to TLF mediated killing. Based on these data, we conclude that the shedding of surface glycoconjugates while transitioning from metacyclic promastigotes to amastigotes results in parasite resistance to TLF mediated lysis. Whether TLF is effective at killing metacyclic promastigotes of other experimentally tractable *Leishmania* sp. such as *L. infantum*, and *L. donovani*, which have slightly different surface glycoconjugate structures is yet to be tested. Our data raise the possibility that TLF can have lytic activity against a broad range of pathogens such as bacteria, viruses, fungi and parasites with surface glycoconjugates that transit through intracellular acidic compartments.

## Introduction

*Leishmania* sp. are intracellular eukaryotic parasites responsible for a spectrum of diseases ranging from mild cutaneous lesions to fatal visceral infections. The parasites are transmitted to humans by the bite of a sand fly vector, wherein the infective motile flagellated, metacyclic form is deposited intradermally into the host. Therein *Leishmania* are taken up by phagocytes and transform into the intracellular amastigote form, which go on to replicate within an acidic parasitophorous vacuole [1]. Although macrophages are the primary host cell for *Leishmania*, parasites can be phagocytosed by other immune cell types including neutrophils and dendritic cells [2-4]. The importance of neutrophils early in infection following the sand fly bite has received increasing attention as a key factor in disease progression, infiltrating the site of the bite as early as 1-hour post-infection and potentially contributing to resolution of the infection [2, 3, 5].

In addition to defenses associated with host cell uptake, *Leishmania* must survive against a variety of extracellular host defenses such as complement [6, 7]. In 2009, our laboratory showed that trypanosome lytic factor (TLF), an extracellular high-density lipoprotein (HDL) complex originally characterized for its ability to kill African trypanosomes, could also ameliorate infection by cutaneous *Leishmania* sp. [8]. TLF is produced only in humans and some higher primates and contains the unique primate proteins haptoglobin-related protein (HPR), and apolipoprotein L-1 (APOL1), the lytic protein [9-14]. Although APOL1 alone is necessary and sufficient for the trypanolytic activity of TLF [13-15], the presence of HPR significantly enhances trypanosome lysis by increasing the uptake of APOL1, via the haptoglobin-hemoglobin receptor (TbHpHbR) [16]. APOL1 then inserts into the parasite endocytic membrane at acidic pH forming a closed ion channel. The acidification is an absolute requirement for the activation/insertion of the APOL1 ion channel [17-19]. The ion channel opens upon neutralization which would occur at the plasma membrane after membrane recycling process, leading to ion flux [19], and consequent ion imbalance resulting in colloid-osmotic lysis.

Human TLF kills the intracellular metacyclic promastigotes of *L. major* and *L. amazonensis* in the phagosomes of macrophages. However, amastigote forms of the parasite are resistant to TLF activity, the basis of which is not well understood [8]. We hypothesize that acidification of phagosomes is essential for TLF-mediated lysis of metacyclic promastigotes within these cells. Following uptake by phagocytes, infective *Leishmania* metacyclic promastigotes can transiently delay phagosomal acidification, mediated primarily by the glycoconjugate lipophosphoglycan (LPG), which is abundant on metacyclic promastigotes but not amastigotes [20-23]. Therefore, infected phagosomes of immune cells may present differing pH environments. Importantly, the lytic component of TLF, APOL1, associates with and forms ion channels in membranes (lipid bilayers) in a pH-dependent manner [18, 19]. The role of phagosomal pH changes in immune cells on the TLF activity against *Leishmania* have not been investigated, nor has the impact of *Leishmania* surface glycoconjugates upon TLF susceptibility.

In this study, we demonstrate that TLF reduces the intradermal parasite burden in transgenic mice producing human or baboon TLF. We find that neutrophils are important for this TLF-mediated protection. Acidification of infected immune cell phagosomes is essential for this TLF-mediated protection. Our *in vitro* data supports this finding, where acidification is essential for TLF-mediated lysis. Amastigotes are resistant to TLF-lysis as reported previously. Finally using glycoconjugate mutants *lpg1*^-^ and *lpg5A*^*-*^ */lpg5B*^*-*^ we show that surface glycoconjugates are required for TLF-interaction and lysis and hypothesize that shedding of the glycoconjugates not only delay acidification of the phagosomes [4, 20-24] but also removes the TLF from the surface of the parasites.

## Results

### Human TLF reduces the parasite burden in a natural intradermal infection by *L. major*

While laboratory mice are good models for Leishmania infection and rodents are a natural zoonotic host, they do not express TLF [8, 9, 13]. In previous work we employed the classic subcutaneous footpad infection model for studying the effect of TLF on *L. major* [8]. Here, we employed the intradermal ear infection model, which more closely mimics the route and numbers of parasites in natural infections [2, 25]. We generated mice that transiently expressed human TLF using hydrodynamic gene delivery (HGD) to introduce the *HPR* and *APOL1* genes, which are secreted and assemble on endogenous murine HDL in the blood, thereby generating TLF [8, 9, 13]. Using this robust expression system, the levels of TLF in the plasma of transfected C57/Bl6 mice after one day are similar to levels in human plasma [8]. These mice were then injected intradermally in the ear with metacyclic promastigotes. Parasites were quantified in the ears 15 days post-infection by qPCR. Consistent with previous studies *in vivo*, parasites numbers were reduced 1000-fold (p= 0.0022) relative to saline injected HGD controls (Fig 1). Thus, we observed TLF-dependent control of *Leishmania*, via two infectious routes, subcutaneous [8] and intradermal herein.

**Figure 1:**
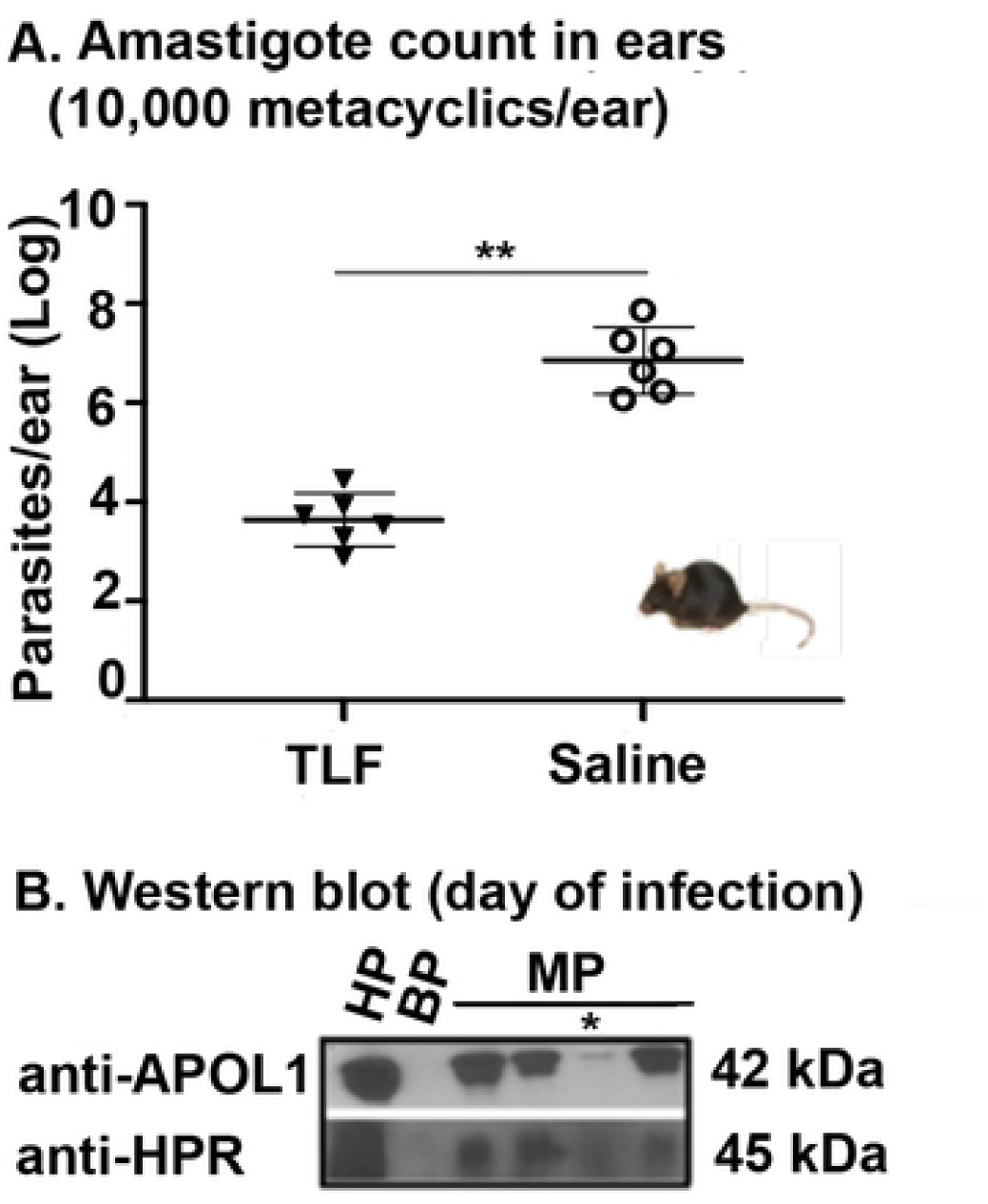
Human TLF reduces the parasite burden in a natural intradermal infection by *L. major*. **A.** Transient transgenic mice were created by injecting 50μg of plasmid DNA containing human *APOL1:HPR* genes in a single plasmid by hydrodynamic gene delivery (HGD) followed by infection by 10,000 metacyclic promastigotes (1 day post-HGD). Parasite burden (amastigotes) was quantified by real time PCR, 15 days post-infection. The data represents mean of one typical experiment that has been repeated three times *p<0.05 Mann-Whitney U test, n= 3 per group. **B.** Western Blots of plasma samples from mice on the day of infection diluted 1:40 (mouse plasma, MP) n=4 were performed to look for the production of APOL1 (42 kDa) and HPR (45 kDa) proteins. * denotes a mouse with failed plasmid injection and hence taken out of the parasite burden analysis. HP (human plasma) diluted 1:40 and BP (bovine plasma) diluted 1:40 were used as positive and negative loading controls.

### Germline targeted transgenic mice expressing baboon TLF ameliorate intradermal infection by *L. major*

While the transient HGD mouse model has many advantages, we found plasmid DNA injection by HGD was highly inflammatory and strongly increased the percentage of CD11b+ (leukocytes) and CD11b+Ly6G+Ly6C^high^ cells (neutrophils) in the blood (S1 Fig), two major cell types known to play important roles in *Leishmania* infection [2, 3]. To explore TLF-*Leishmania* interactions in a less perturbative setting *in vivo*, we took advantage of a recently produced germline transgenic murine model that produces baboon TLF (we do not have germline transgenic human TLF mice). As expected, the APOL1 and HPR in these mice can be detected in lipoprotein complexes upon density gradient centrifugation and, as in human plasma, both proteins elute together with structural protein APOA1 in ∼500kDa HDL fractions after size-exclusion chromatography (fractions 11-13), suggesting that a complete TLF complex was reconstituted in these mice (Fig 2A). Hence, we expect the TLF of these mice to restrict *Leishmania* like human TLF in the transiently transfected mice (Fig 1). We infected the germline transgenic mice with 10^4^ metacyclic promastigotes. We did not see a difference in the number of parasites (quantified by qPCR) between TLF expressing and the control mice 15 days post-infection. However, we observed a ten-fold reduction in parasite burden at an infective dose of 10^3^ metacyclic promastigotes (p=0.1303) and 100 metacyclic promastigotes (p<0.0001) (Figs 2B-D). Our results show that the outcome of TLF activity against *L. major* is governed in part by the infective dose of parasites. We also observed that the magnitude of any TLF-mediated protection also depends on the host’s circulating TLF/HDL concentration. Homozygous mice with two copies of *APOL1* and *HPR* produce twice as much proteins as heterozygous mice (S2A Fig). Accordingly, homozygous mice show a ten-fold reduction (p=0.1303) in parasite burden compared to the heterozygous mice that show only a five-fold reduction (p=0.5722) (Fig 2C) suggesting that TLF lytic activity is dependent on the circulating levels of TLF proteins APOL1 and HPR (S2B Fig). Our results suggest that both human TLF and baboon TLF can reduce the parasite burden in *L. major* infection after intradermal delivery of the parasites. The final outcome of the disease depends on both the initial dose of infecting parasites, the concentration of circulating TLF in the plasma and the immune cells recruited to the site.

**Figure 2:**
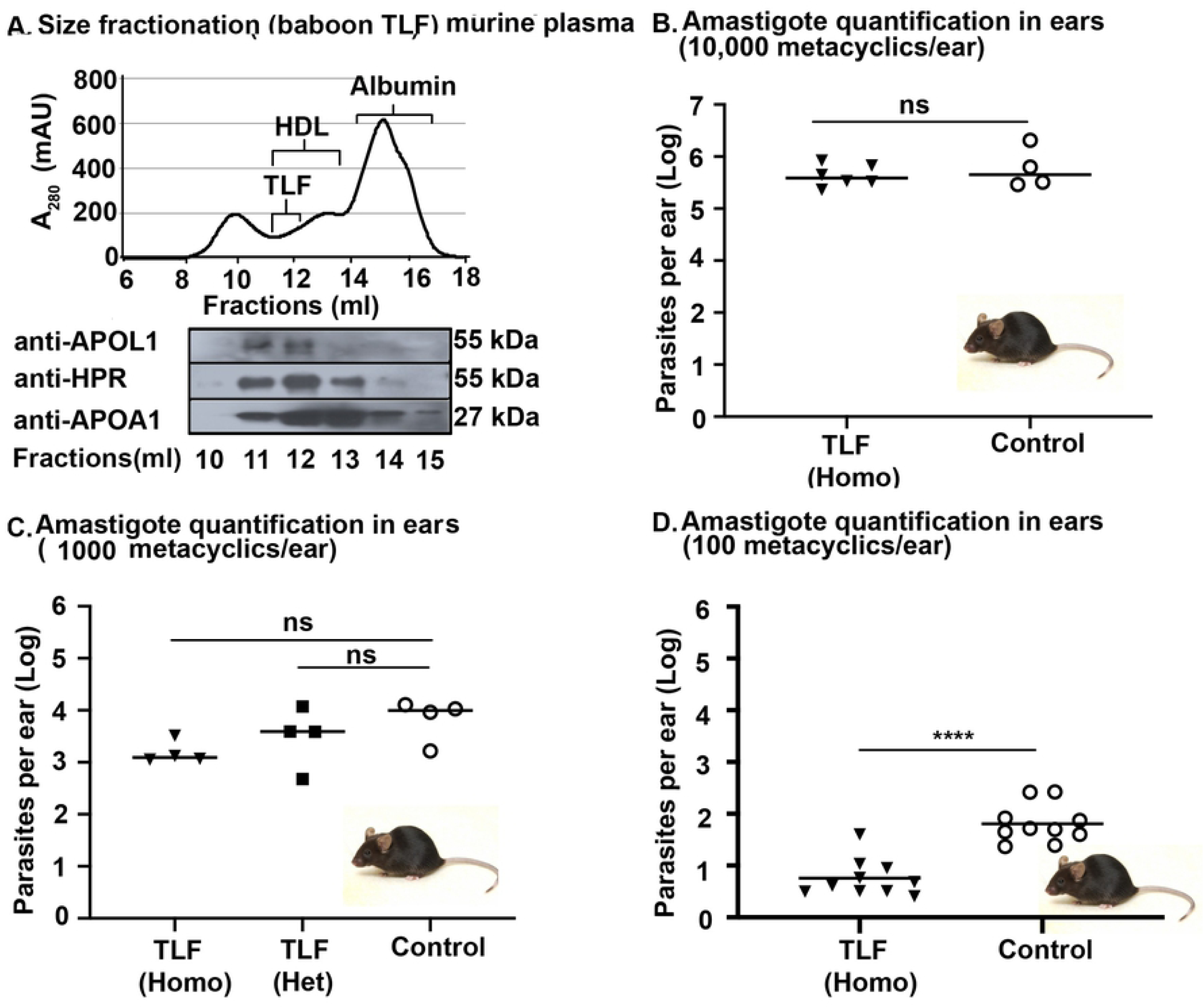
Germline targeted transgenic mice expressing baboon TLF ameliorate intradermal infection by *L. major*. **A.** Plasma from germline transgenic mice expressing baboon *APOL1* and *HPR* separated by size exclusion chromatography showing the protein absorbance profile (280 nm) and the western blot performed of the fractions (11–15) for three TLF proteins-APOL1, HPR and APOA1. Fractions 11–14 contain HDL. The proteins (APOL1, HPR and APOA1) were also detected in the lipoprotein layer collected from density gradient centrifugation (Lipo) from transgenic mice **B–D.** Number of parasites in germline transgenic mice (Homo = homozygotes, Het = Heterozygotes) ears quantified by real time PCR 15 days post-infection with **B.** 10,000 metacyclic promastigotes n=6 (TLF), n=4 (control). **C.** 1000 metacyclic promastigotes, n=4 per group. **D.** 100 metacyclic promastigotes, n=10 per group. The data represent Mean ± SD of one typical experiment that has been repeated three times. *p<0.05, ns-non-significant, Mann-Whitney U test was used for statistical analysis.

### Neutrophils are required for TLF-mediated protection against *L. major in vivo*

Neutrophils are the first cells recruited and infected by metacyclic promastigotes in an intradermal infection by *L. major* [2]. We depleted the neutrophils from the transgenic mice expressing baboon TLF (homozygotes) and control (wild-type) mice by injecting anti-mouse Ly6G 1A8 antibodies. Isotype matched IgG antibody was injected as a control. After 24 hours, we observed a 50% reduction of neutrophil frequency in the blood (Fig 3A). At this time, we infected mice with 100 metacyclic promastigotes per ear. At 10 h post-infection (peak time of neutrophil recruitment), we also observed a reduction of neutrophils in the ear (site of infection) of mice injected with anti-mouse Ly6G 1A8 antibodies (Figs 3C and S3 Fig). The TLF mice with depleted neutrophils had a significantly higher burden of amastigotes in their ears (site of infection) compared to the isotype control mice, at 15 days post-infection (p=0.0271). This was contrary to the result for wild-type mice wherein depleting neutrophils resulted in a significant reduction (p=0.0201) in parasite burden in their ears compared to the mice that received the isotype control antibody (Fig 3B). The data suggest that neutrophils promote infection by acting as a safe harbor for the parasites in the absence of TLF, as has been previously reported [2-4]. However, in the presence of TLF as in all humans, neutrophils are not a safe harbor for the parasites and are instead required for efficient control of the infection. These data reveal a dynamic interaction between TLF, metacyclic promastigotes of *L. major*, and neutrophils providing innate immunity against intradermal parasites. Previously, we had observed a colocalization of TLF and *L. major* metacyclic promastigotes in the parasitophorous vacuoles of macrophages [8]. Therefore, we hypothesize that metacyclic promastigotes in our TLF-expressing mice encounter TLF in the acidic endosomes of phagocytes upon intradermal infection, where they are subjected to the lytic activity of TLF.

### Acidic pH is essential for TLF-mediated lysis of *L. major*

To test if metacyclic promastigotes are lysed in the acidic parasitophorous vacuoles of phagocytes, we infected macrophages with metacyclic promastigotes, and incubated the cells with TLF or bovine HDL (bHDL; cattle do not make TLF) in the presence of ammonium chloride (NH_4_Cl, S4 Fig), to neutralize the endocytic compartment of the macrophages or absence of NH_4_Cl. Addition of TLF in the absence of NH_4_Cl significantly decreases the parasite number within the macrophages as previously observed (p= 0.0052) [8]. However, in the infected macrophages subjected to neutralization treatment (NH_4_Cl), TLF had no effect on the parasite number. Additionally, treatment of infected macrophages with bHDL with or without NH_4_Cl had no effect on the parasite number (Fig 4A). These data show that neutralization of the acidic compartments within macrophages inhibits the lytic activity of TLF against *L. major.* Therefore, the acidic pH of macrophage phagocytic compartments is essential for TLF activity against *Leishmania*. Next, we tested the effect of TLF on *Leishmania* at various pH conditions *in vitro*, to test the pH conditions that broadly occur in phagocytes. As we cannot obtain axenic amastigotes from *L. major*, we used *L. amazonensis*, another TLF susceptible cutaneous strain [8]. There was no killing of the metacyclic promastigotes at any pH by the bHDL control (Fig 4B). The data are expressed as a ratio of surviving parasites post TLF-treatment to bHDL-treatment. A ratio of one would indicate equivalent survival in both treatments, while less than one would indicate increased parasite killing by TLF-treatment. Metacyclic promastigotes subjected to increased acidification (Fig 4B, IA, pH5.6-4.5) in the presence of TLF were completely lysed in two hours (p<0.0001). We also observed a significant reduction (p=0.0036) in parasite number when metacyclic promastigotes were incubated with TLF and subjected to acidification (pH 5.6 for 1 hour) followed by neutralization (pH 7 for 1 hour) (Fig 4B, AN). This reduction in parasite number was not observed in metacyclic promastigotes incubated with TLF at neutral pH for 2 hours (Fig 4B, N). Therefore, the pH of the milieu governs the susceptibility of metacyclic promastigotes to TLF-mediated lysis. Acidification is required for the lysis of metacyclic promastigotes by TLF, which occurs upon exposure to acidic pH conditions within the phagosomes of immune cells [8, 20-22 24]. Consistent with our previous results, we did not observe lysis of axenic amastigotes by TLF suggesting that axenic amastigotes are resistant to TLF activity (Fig 4C). Therefore, TLF-mediated protection is restricted to the early stage of infection when the parasite exists as metacyclic promastigotes. The fact that metacyclic promastigotes are lysed by TLF while amastigotes are resistant (Figs 4B & C) suggests that there could be a specific interaction between TLF and the surface of metacyclic promastigotes. Therefore, we analyzed the binding between TLF and both metacyclic promastigotes and axenic amastigotes. We observed that TLF and bHDL interacts with metacyclic promastigotes but not with the resistant amastigotes (Figs 4D & S5 Fig) suggesting that TLF-mediated lysis could be affected by the differences between the cellular surfaces of the two life-cycle stages.

**Figure 3:**
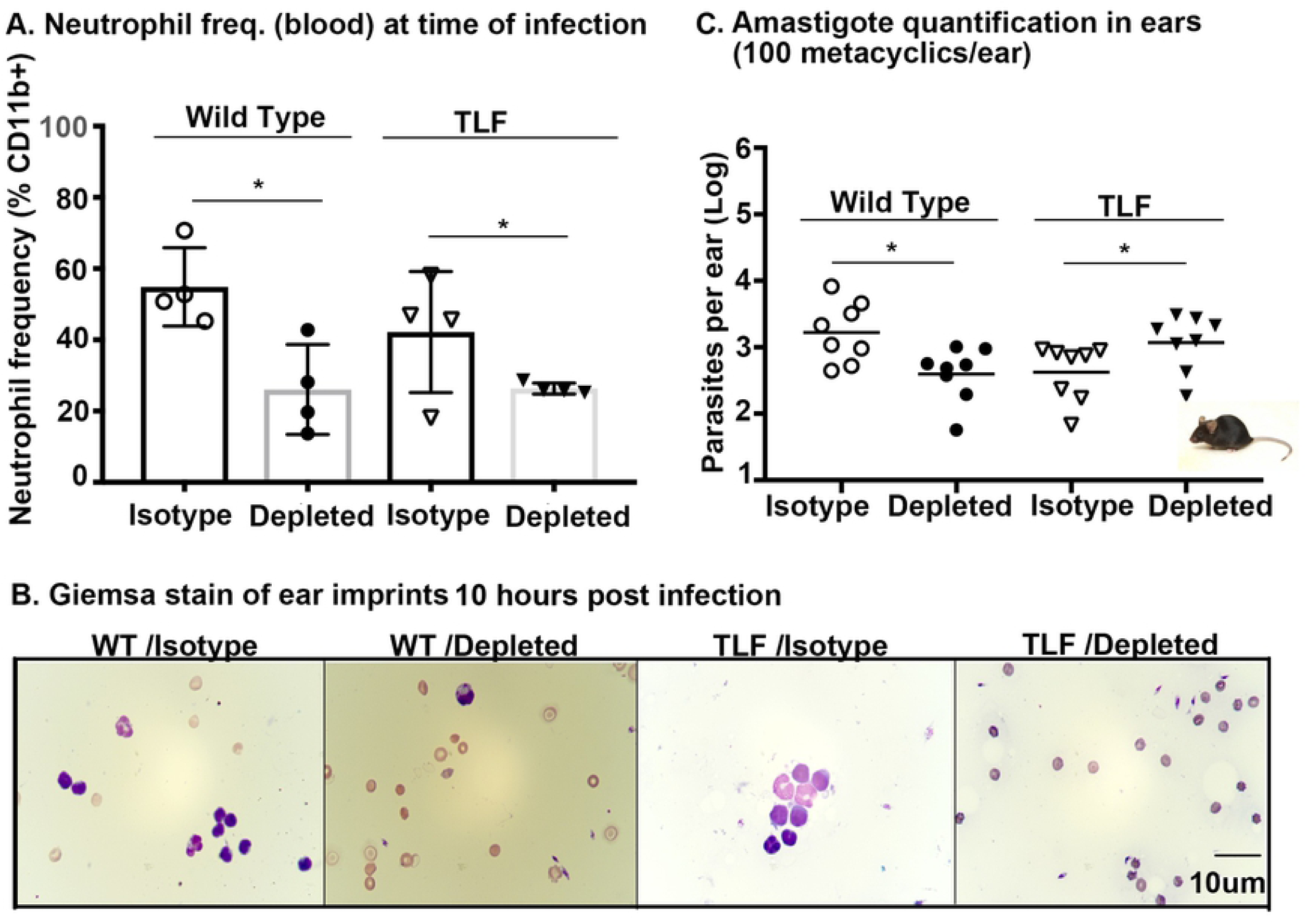
Neutrophils are required for TLF-mediated protection against *L. major in vivo*. Neutrophils were depleted in mice using 1 mg anti-mouse Ly6G clone 1A8 antibody (depleted) or not depleted using an isotype matched IgG2A antibody (isotype). **A.** Neutrophil frequencies were assayed 24 hours post-depletion (time of infection) by collecting blood from the tail, staining for surface markers and analyzing by flow cytometry. The bar graph depicts the neutrophil frequency (CD11b^+^Ly6G^+^Ly6C^+^). **B.** The number of parasites quantified by real time PCR in mice ear harvested 15 days post-infection with 100 metacyclic promastigotes. The data represents Mean ± SD of one typical experiment that has been repeated twice. * *p*<0.05, Mann-Whitney U test, n=4 per group. **C.** Giemsa stains of the ear imprints collected 10 hours post-infection from wild-type mice and TLF mice with (isotype) or without neutrophils (depleted) visualized using Nikon Eclipse 50i microscope under 100X oil immersion. The data is representative of multiple fields visualized that are repeated twice.

**Figure 4:**
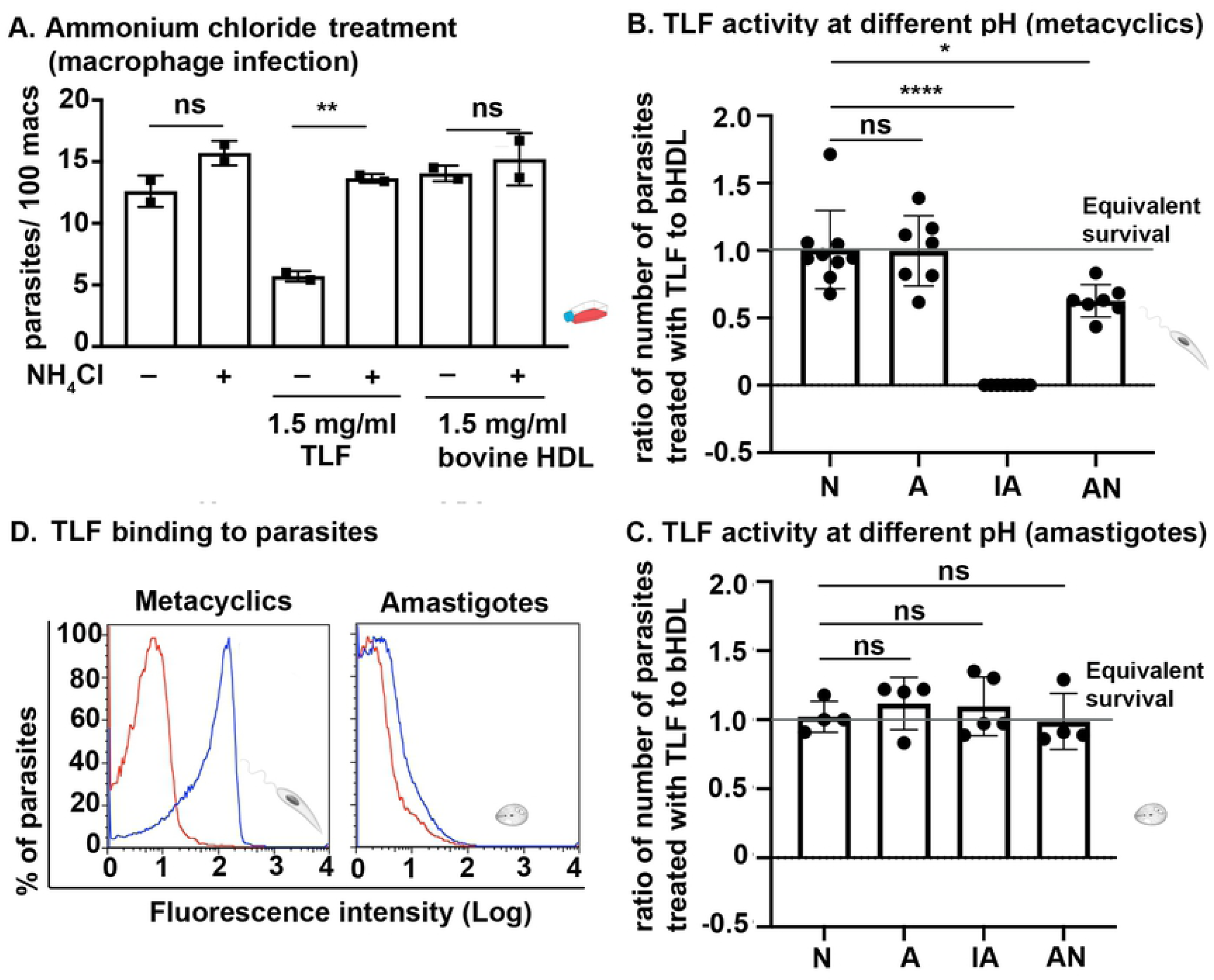
Acidic pH is essential for TLF-mediated lysis of *L. major*. **A.** Bone marrow derived macrophages from BALB/c mice were infected with *L. major* metacyclics at a multiplicity of infection of 3:1, for 2 hours, before treatment with NH_4_Cl (10 mM, where indicated) for 30 min followed by the addition of TLF or bovine HDL (1.5 mg/ml) for 24 hours. 10 mM NH_4_Cl was maintained throughout the 24 hours, to maintain neutralization of macrophage endocytic compartments. The data represent the mean ± SD of duplicate cultures of one typical experiment repeated twice. ** *p*<0.001 TLF compared to bovine HDL, ANOVA test. **B.** Metacyclic promastigotes or **C.** Amastigotes of *L. amazonensis* were treated with TLF (1.5 mg/ml) or bovine HDL (1.5 mg/ml) and incubated in different conditions, Neutral, N-pH 7.4 for 2 h, Acidic, A-pH 5.6 for 2 hours, Increased acidification, IA-pH 5.6 for 1 hour followed by pH 4.5 for 1 hour, Acid and Neutral AN-pH 5.6 for 1hour followed by pH 7.4 for 1hour. Parasites were microscopically counted using a hemocytometer. The data represent Mean ± Standard Deviation (SD) of three experiments that have been combined. **** *p*<0.0001, ns-non-significant, ANOVA test was used for statistical analysis. **D.** *L. amazonensis* metacyclic promastigotes or axenic amastigote forms (1×10^6^/ml) were treated with 10 µg/ml of DyLight-488 labelled TLF (blue) or not (red) for 30 min on ice. Fluorescence intensity was quantified by flow cytometry. The data represent one typical experiment repeated three times.

### *S*urface glycoconjugate mutants of *L. major* are less susceptible to TLF-mediated lysis than wild-type parasites

Amastigotes, which are the pathogenic forms of *Leishmania*, have distinct surface glycoconjugate components compared to the infective metacyclic stage [23]. Therefore, we hypothesized that surface glycoconjugates attached to the plasma membrane of the parasites play an important role in TLF-binding and hence TLF-susceptibility. To evaluate the role of the surface glycoconjugates in differential susceptibility to TLF-mediated lysis of metacyclics, we utilized mutant lines of *L. major* parasites deficient in the synthesis of specific surface structures. The *lpg1*^-^ mutants lack the lipophosphoglycan (LPG) core galactofuranosyl transferase LPG1 and are specifically deficient in LPG synthesis [26]. The *lpg5A*^*-*^*/lpg5B*^*-*^ double mutants cannot import UDP-Galactose (UDP-Gal) into the Golgi and are therefore defective in the synthesis of multiple glycoconjugates including LPG and proteophosphoglycan (PPG) [27]. We tested the protective effect of TLF against these mutants by infecting macrophages with *lpg1*^-^ or *lpg5A*^*-*^*/lpg5B*^*-*^ metacyclic promastigotes in the presence of TLF or bHDL. Wild-type *L. major* parasites were used as a control. The metacyclic promastigotes of *lpg1*^*-*^ and *lpg5A*^*-*^*/lpg5B*^*-*^ mutants were less susceptible to TLF-mediated lysis compared to the wild-type *L. major* strain (Fig 5A). Despite being less susceptible to TLF, *lpg1*^*-*^ and *lpg5A*^*-*^ */lpg5B*^*-*^ mutants were still able to bind labelled TLF (Fig 5B). Complemented cells expressing *LPG1* (Fig 5A) demonstrate WT phenotypes but TLF susceptibility was only partially restored in the complemented line *lpg5A*^*-*^*/lpg5B*^*-*^*/+LPG5A+LPG5B* (Fig 5A). To confirm that the complemented lines were producing glycoconjugates, we screened the level of phosphoglycans in these mutants and compared it to wild-type *L. major*. Although the level of phosphoglycans was not restored to the wild-type parasites’ level, the complemented lines produce phosphoglycans (Fig 5C). Despite the low level of phosphoglycan production, TLF susceptibility was restored in the complemented lines. Our results support the hypothesis that the surface glycoconjugates of metacyclic promastigotes contribute to the susceptibility of TLF-mediated lysis in macrophages.

**Figure 5:**
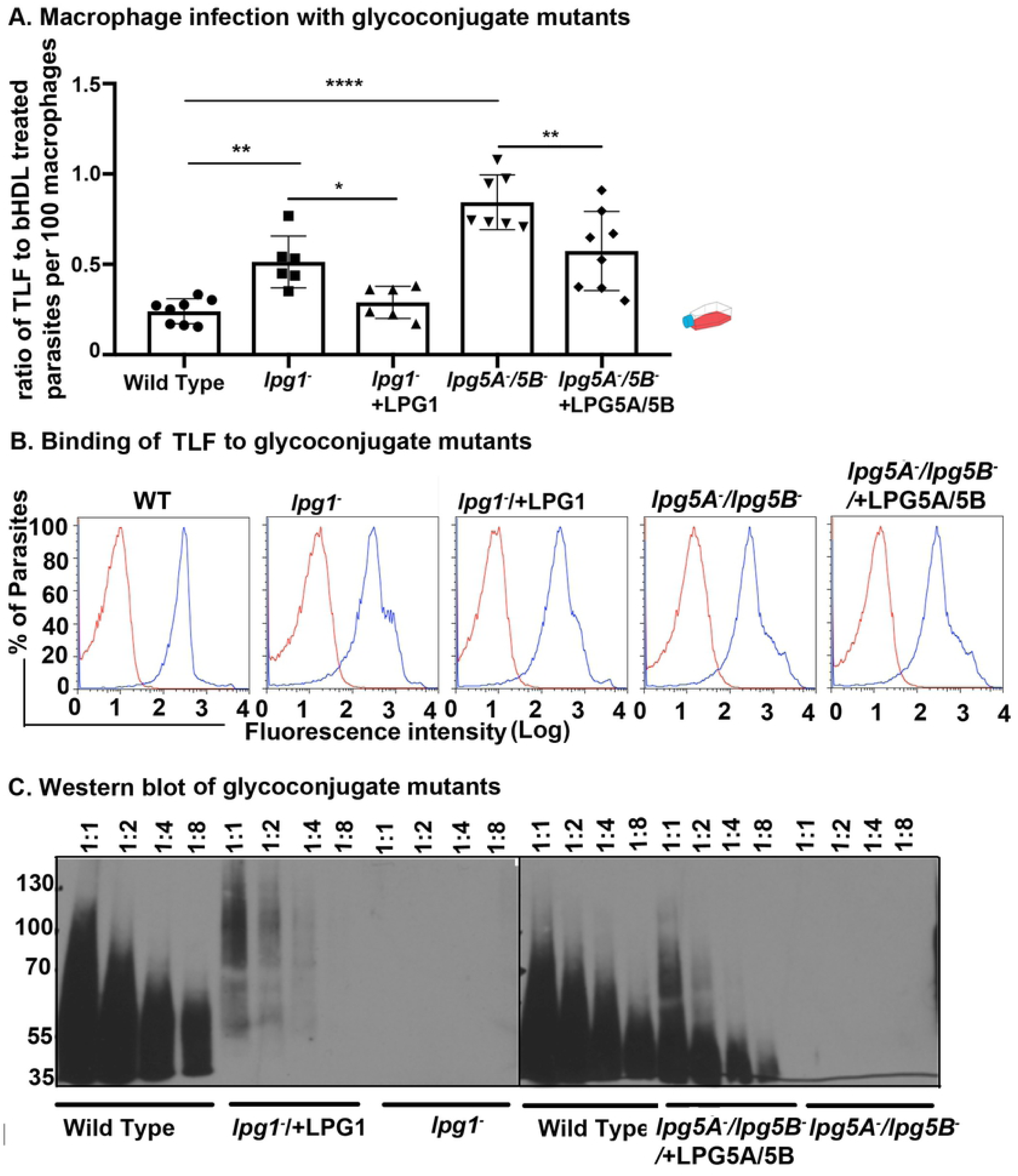
*S*urface glycoconjugate mutants of *L. major* are less susceptible to TLF-mediated lysis than wild-type parasites. Metacyclic promastigotes of *L. major* (WT), or glycoconjugate mutants (*lpg1-* and *lpg5A*^*-*^ */lpg5B*^*-*^), or complemented lines (l*pg1*^*-/-*^+LPG1 and *lpg5A*^*-*^*/lpg5B*^*-*^+LPG5A/5B) were **A.** used to infect macrophages in the presence of TLF or bovine HDL (1.5 mg/ml). Parasites were counted in macrophages at 24 hours post-infection by staining with DAPI. The y-axis represents the ratio of parasites per 100 macrophages resulting from TLF treatment compared to bHDL treatment. The data are representative of three experiments combined, * *p*< 0.05, ** *p*<0.001, and **** *p*<0.0001. ANOVA test was used for statistical analysis. **B.** co-incubated with 10 µg/ml of DyLight-488 labelled TLF (blue) at 4°C for 30 min. The fluorescence was measured using flow cytometry where untreated parasites (red) were used as control. **C.** were lysed (2×10^7^) and the extracts were then serially diluted to compare the expression of phosphoglycan (PG) proteins, which were probed using anti-phosphoglycan antibody WIC 79.3.

## Discussion

TLF is an innate immune complex in humans and some non-human primates that was originally discovered due to its protective effect against African trypanosomes. Previously, we have shown that human TLF ameliorates the infection caused by other intracellular kinetoplastid parasites *L. major* and *L. amazonensis* both in macrophages and in a subcutaneous model of infection in transgenic mice [8]. In the subcutaneous model, macrophages are the predominant cells that phagocytose the parasites early in the infection [3]. Therefore, it can be inferred that TLF lyses metacyclic promastigotes of *L. major* in the macrophages *in vivo*. However, the role that TLF plays during natural intradermal infections by *Leishmania* is not known. In this study, we found that TLF can ameliorate infection by cutaneous *L. major in vivo* when mice are injected intradermally with metacyclic promastigotes (Fig 1). Therefore, TLF-mediated innate immunity could be important during natural sand fly infections in humans.

TLF activity has been detected in a few primates such as humans, gorillas, baboons, mandrills, and sooty mangabeys [9, 10, 28]. While hominoid (humans and gorillas) TLF only protects against animal-infective trypanosomes, old-world monkeys (OWMs) have TLFs that confer resistance against both human and animal-infective trypanosomes. This differential activity is attributed to differences in the APOL1 sequence which is approximately 60% identical between human and baboon [9] showing the evolutionary divergence of the protein in the two species. Here, we found that both human and baboon TLFs protect against *L. major* (Figs 1 & 2). However, transiently transgenic mice (generated by HGD) producing human TLF were able to reduce the parasites when inoculated with a higher infective dose than germline targeted transgenic mice producing baboon TLF (Figs 1 & 2B). This difference in killing activity between transiently transgenic HGD mice and germline baboon TLF-producing transgenic mice could be due to: 1) A mechanistic difference in activity between human and baboon TLF which should be further studied in future; 2) Increase in immune cells induced by HGD (S3 Fig); 3) Differences in the concentrations of circulating TLF (S2B Fig). Our data provides strong evidence to support that the differential killing of *L. major* observed in transiently transgenic mice and germline transgenic mice is due to the difference in the level of circulating TLF in these mice (Figs 1 & 2 and S2B Fig). The homozygous germline transgenic mice produce twice as much TLF proteins (APOL1-1:40 dilution; HPR 1:160 dilution) compared to heterozygous mice (APOL1-1:20 dilution; HPR-1:80 dilution). As expected, homozygous germline transgenic mice show a 10-fold reduction in parasite burden compared to 5 fold in heterozygous mice (Fig 2C). This data further supports the hypothesis that the difference in circulating levels of TLF proteins APOL1 and HPR can lead to variation in TLF-mediated protection. In addition, we found that the TLF-mediated reduction in the parasite burden in the mice was also dependent on the initial infective dose (Fig 2). In a natural infection, the disease outcome possibly depends on the number of parasites that the sand fly injects as well as the circulating level of HDLs and TLFs in each individual. Therefore, low HDL level at the time of infection possibly exacerbates *L. major* infection. In fact, infection by visceral leishmania strains has been associated with a reduction in HDL compared to the healthy control subjects in human studies [29, 30].

Following infection, *L. major* recruits phagocytic immune cells to the site of infection. Although the number and type of these phagocytes are variable depending on the route of infection, neutrophils represent the predominant cells that phagocytose the parasites early post-infection irrespective of the infective route [3]. Neutrophil depletion experiments in mice have shown controversial results in terms of whether the role of neutrophils is primarily protective or deleterious for *Leishmania* [2, 3, 5, 21, 31, 32]. Notably, these early experiments were performed in wild-type mice that do not produce TLF. We found that depleting neutrophils prior to infection (reduced neutrophil recruitment at early hours post-infection) leads to a reduction in parasite burden in wild-type mice not producing TLF when intradermally infected with *L. major* (Fig 3B). This was in agreement with previous results [2, 3, 5]. On the contrary, we observed a significant increase in parasite burden in TLF mice depleted of neutrophils compared to the control mice with TLF and neutrophils (Fig 3B). This result supports our hypothesis that TLF kills metacyclic promastigotes of *L. major* in neutrophils. This opens the possibility that TLF can act on *L. major* within the acidic phagosomes of other immune cells, which should be investigated in future studies.

Phagosomes of immune cells are subject to modulation by parasites where parasite LPG inhibits the acidification of resident phagosomes within the first 2 hours post-infection [4, 20-22, 24]. Therefore, TLF activity against parasites within acidic phagosomes of immune cells must occur within the first 1-2 hours post-infection. Our data shows that acidification of phagosomes is essential for TLF-mediated killing of *L. major*, which can be inhibited by neutralizing the phagosomes of macrophage with the weak base ammonium chloride (Figs 4A & S4 Fig). To focus purely on pH-dependent activity of TLF against *Leishmania*, we analyzed parasite susceptibility to TLF at different pH conditions *in vitro*. We observed 100% lysis when parasites were subjected to increasing acidification (IA) from pH 5.6 to pH 4.5 to mimic maturing phagosomes. We do not know the mechanism of lysis in metacyclic promastigotes in increasing acidification conditions. We tried to investigate this using artificial bilayers using recombinant APOL1. However, artificial bilayers were unstable at pH below 5 in the presence of APOL1. We interpret these data as the possible increase in insertion of APOL1 into membrane bilayer at acidic pH that leads to membrane disruption.

There is partial but significant lysis of parasites upon acidification at pH 5.6 followed by neutralization (AN) that can occur in phagosomes in the presence of phosphoglycan such as LPG (Figs 4B & C) [4, 20-22, 24]. Nonetheless, there are always some parasites that survive and establish an infection in the presence of TLF in both *ex vivo* macrophage infection and *in vivo* mouse infections (Figs 1, 2 & 4A). Therefore, it is highly likely that infected immune cell phagosomes follow the acidification and then neutralization route and not the gradual acidification that leads to 100% lysis of parasites. This is analogous to the APOL1-mediated lysis model as observed in artificial bilayers, where recombinant APOL1 inserts into the bilayer and forms a closed-channel at acidic pH. This closed state channel subsequently opens upon neutralization allowing movement of cations [19]. We hypothesize that the APOL1 protein of the TLF complex forms cation channels in the parasite plasma membrane that eventually leads to membrane depolarization and lysis, which will be further investigated.

There is a fundamental difference in the mechanism of lysis of *Leishmania* by TLF compared to African trypanosomes. In African Trypanosomes, when TLF or APOL1 is co-incubated with parasites in a neutral pH milieu (blood and extracellular fluid), the parasites endocytose TLF/APOL1, which inserts into the membrane of acidic endo/lysosomes to form a closed ion channel. The pH-gated ion channel opens upon neutralization, which occurs when the endo/lysosomal membrane is recycled to the plasma membrane leading to ionic imbalance, osmotic influx of water and lysis of the parasites [18, 19, 33]. In contrast, incubation of TLF with *L. major* metacyclic promastigotes in a neutral pH milieu (plasma and extracellular fluid) does not kill the parasites (Figs 4B) & [8], and we have been unable to obtain any evidence suggesting that *Leishmania* parasites endocytose detectable TLF. However, increasing acidification (pH 5.6 to pH 4.5) or acidification followed by neutralization (pH 5.6 to pH 7) of the milieu results in parasite lysis (Fig 4). Therefore, the lysis of *L. major* metacyclic promastigotes by TLF occurs within phagosomes of immune cells, forming ion channels in the surface of the parasites. This is further supported by the fact that amastigotes with a different glycoconjugate composition, compared to metacyclic promastigotes, do not bind TLF and are resistant to TLF lysis (Fig 4C). Moreover, mutant parasites deficient in surface glycoconjugates are also resistant to TLF-mediated lysis. These data provide strong evidence to support the hypothesis that TLF binds to the surface of metacyclic promastigotes, allowing APOL1 to form pH-gated ion channels in their plasma membrane leading to their colloidal osmotic lysis.

Denuding parasites of surface glycoconjugates of *Leishmania* sp. renders them resistant to TLF lysis (Fig5A) suggesting that TLF interacts with surface glycoconjugates of the metacyclic promastigotes. Metacyclic promastigotes shed their surface glycoconjugates LPG into phagosomal membranes, which interferes with phagosomal maturation [4, 20-22, 24]. Therefore, shedding of surface glycoconjugates may be one of the mechanisms used by parasites to evade TLF-mediated lysis within phagosomes. Our binding experiments show that TLF binds to surface glycoconjugates deficient parasites similar to wild-type metacyclic promastigotes (Fig 5B). However, this binding of TLF to the glycoconjugate deficient metacyclic promastigotes is not sufficient to lyse the parasites (Fig5A). Infact, bHDL that does not kill parasites also shows binding to the metacyclic promastigotes of *Leishmania* (S5 Fig) suggesting that specific interaction between TLF and surface glycoconjugates is required for TLF-susceptibility. The fluorescence intensity of TLF bound parasites is greater than the bHDL parasites, which is possibly due to the difference in labelling of the HDL by Dylight-488. In our hands, labelling of bHDL was always less efficient than TLF possibly due to the difference in the protein composition between two complexes. The TLF complexes are characterized by presence of 60% proteins, 40% lipids (density-1.21-1.24) compared to other HDL complexes 50% proteins, 50% lipid (density-1.063-1.21 g/ml) [33]. Our data show that although other HDL complexes bind the parasite surface, only TLF-binding results in lysis.

Our results show that TLF has evolved in primates as an innate immune factor that kills extracellular African Trypanosomes and intracellular *Leishmania* sp.. The parasites have evolved to overcome the TLF activity and hence can eventually survive, divide, and proliferate to cause disease pathogenesis. The African Trypanosomes strains *Trypanosoma b. rhodesiense and T. b. gambiense* have evolved to partially evade TLF activity and cause human African trypanosomiasis [35]. Likewise, *Leishmania* sp. have evolved to transition into amastigotes with different surface glycoconjugates that can evade TLF activity. Therefore, presence of parasite surface glycoconjugates determines the susceptibility of metacyclic promastigotes to TLF-mediated lysis. Our data infer that innate immunity afforded by TLF may be effective against a broader range of intracellular pathogens coated with diverse glycoconjugate moieties.

The final outcome of disease depends on the initial dose of pathogens and the concentration of circulating TLF complexes in addition to the type of interacting immune cells.

## Materials and Methods

### Parasites and media

*L. major* strain Friedlin V1 (MHOM/JL/80/Friedlin) or *L. amazonensis* (IFLA/BR/67/PH8), *lpg1*^*-/-*^, *lpg1*^*-/-*^/*+LPG1, lpg5A*^*-*^*/lpg5B*^*-*^*/+LPG5A+LPG5B* promastigotes were grown in M199 medium (Gibco) [26, 27]. The parasites were grown to stationary phase culture. Infective-stage metacyclic promastigotes were isolated by density centrifugation on a Ficoll (Sigma, F8016) gradient from an 8–10 day culture [36].

### Murine transfection and infection

Transient transgenic mice were created by injecting 50 µg of plasmid DNA (*APOL1:HPR*) by hydrodynamic gene delivery (HGD) as described previously [8, 9, 13, 37]. One day post-HGD, plasma was collected by tail bleed to analyze the expression level of both genes by western blot using a rabbit polyclonal anti-APOLI antibody (ProteinTech 16139-1-AP - 1:10,000) or a rabbit polyclonal anti-HP antibody (Sigma H8636 - 1:10,000) as the primary antibody and TrueBlot anti-rabbit secondary antibody (Rockland Antibodies 18-8816-33 - 1:5000). Transiently transfected mice were intradermally infected on the same day (1 day post-HGD transfection) with 10^4^ Ficoll (Sigma) purified metacyclic promastigotes in the ear. This dose was chosen based on previous studies showing that a sand fly injects a wide range of parasites intradermally during its blood meal that ranges from 10–10^5^ metacyclic promastigotes per bite [25]. The sand fly infection dose follows a bimodal distribution with a high median dose ranging from 10^3^–10^4^ parasites and the lower median dose from 100–600 parasites [25]. Parasite dose ranging from 100 to 10^4^ were intradermally injected into mice as indicated in the figures. For parasite quantification, ears were harvested on day 15 post-infection and quantified by qPCR as described below.

Germline transgenic mice were generated by targeted recombination into the ROSA 26 locus. The baboon *APOL1* gene is under the control of a ubiquitin promoter and the baboon *HPR* gene is under the control of an albumin promoter. The ROSA26 locus is used in mice for constitutive expression of genes. Details of the mice will be published elsewhere. APOL1 is expressed in various tissues in primates [37]. Expression of HPR is however, limited to the liver [38]. Our germline murine model thus resembles the expression of these genes in the primates. All animal experiments were approved by the Hunter College Institutional Animal Care and Use Committee (IACUC), which has current approved animal welfare assurance agreement number of D16-00413 (A3705-01) from National Institute of Health, office of Laboratory animal welfare.

### APOL1 and HPR produced in plasma from germline transgenic mice

Plasma was collected by tail bleeding from germline transgenic homozygous and heterozygous mice expressing *APOL1* and *HPR.* For comparing APOL1 and HPR production, we serially diluted the plasma and separated by non-reducing SDS PAGE gel as described above. The proteins were then probed for baboon APOL1 using anti-baboon APOL1 antibody (rabbit anti-sera raised against the following peptide: CSVEERARVVEMERVAESRTTEVIRGAKIVDK, AnaSpec Incorporated, 1:1000) and anti-Hp antibody (H8636 - 1:10,000). Baboon HPR has two predicted glycosylation sites and hence runs higher (∼ 50 kDa) than the expected protein size [10]. Baboon APOL1 runs higher (∼55 kDa) than its expected size of 42 kDa.

### Parasite number determination by qPCR

For parasite number determination by qPCR, ears were harvested 15 days post-infection, digested in trizol (ZYMO Research) and DNA was isolated. Parasite DNA was quantified using previously described primers that amplify the kinetoplastid DNA (kDNA): forward – 5’-CCTATTTTACACCAACCCCCAGT-3’ [JW11]; reverse - 5’-GGGTAGGGGCGTTC TGCGAAA-3’ [JW12] [40, 41]. The murine housekeeping genes used were beta-catenin and GAPDH amplified using previously described primers: b-cat-F 5’-CTTGGCTGAACCATCAC-3’; b-cat-R 5’-GGTCCTCATCGTTAGCA-3’; GAPDH-F 5’-TTTGATGTTAGTGGGGTCTCG-3’; GAPDH-R 5’-AGCTTGTCATCAACGGGAAG-3’ [38, 39]. PCR was performed using the Applied Biosystems ViiA Real Time PCR System in a 20 µl reaction mix with 10 µl SYBR Green master mix (Applied Biosystems), 1 µmol each of the forward and reverse primers and 50ng of the template. The PCR was run for 40 cycles at 95°C for 10 sec, 60°C for 15 sec and 72°C for 30 sec. Parasite Ct were normalized against mouse GAPDH Ct and the numbers of parasites were quantified using the standard curve of mouse ear tissues with known numbers of *Leishmania* parasites (limit of linear detection ranges from 10^2^ to 10^6^ parasites per ear).

### Neutrophil depletion and infection

Neutrophils were depleted from targeted transgenic mice expressing baboon TLF (APOL1 and HPR homozygotes) and wild-type C57BL/6NTac (Taconic) mice not expressing TLF by injecting anti-mouse Ly6G, clone 1A8 antibodies intraperitoneally, (BioXcell, BP0075-1, 1 mg). Isotype matched IgG2a, clone 2A3 (BioXcell, BP0089, 1mg) antibody was injected as a control. Blood was collected 24 h later and stained with PE Rat Anti-Mouse CD11b-clone M1/70, FITC Rat Anti-Mouse Ly-6G and Ly-6C clone RB6-8C5, APC Rat Anti-Mouse Ly-6C-Clone AL-21 (BD Pharmingen) for 30 minutes in the dark. The samples were then treated to BD FACS™ lysing solution for 10 min at room temperature followed by two washes. Flow cytometry was performed with a Becton Dickinson FACSCalibur system and BD FACScan.

After blood collection mice were infected with metacyclic promastigotes (100 for ear infection kinetics and 10^6^ for neutrophil analysis in ears). To check neutrophil recruitment, ears were harvested from mice 10 hours post-infection (peak neutrophil recruitment time) and imprinted onto slides followed by methanol fixation and stained with Giemsa. To check by flow cytometry, the ears were digested with 0.2mg/ml Liberase (Roche; 05401020001) for 2 hours at 37°C. Cells were placed on ice and stained as described above for blood and assayed by flow cytometry.

For quantification of parasites in ears, ears were harvested 15 days post infection and processed as described above.

### Purification of TLF, bovine HDL and mouse HDL

The HDL from normal human plasma was purified by density gradient centrifugation as described elsewhere [15, 42]. The plasma density was adjusted to 1.25 g/ml using potassium bromide (KBr) and centrifuged at 49,000 rpm (NVTi 65; Beckman) for 16 hours at 10°C. The top lipoprotein fraction was collected followed by density adjustment to 1.3 g/ml with KBr. Then 4 ml of the adjusted lipoprotein was layered under 8 ml of 0.9% NaCl and centrifuged at 49,000 rpm for 4 hours at 10°C (NVTi 65 rotor; Beckman). The HDL (lower band) was then harvested and dialyzed in Tris-buffered saline [TBS; 50 mM Tris-HCl, 150 mM NaCl (pH 7.5)] at 4°C and concentrated by ultrafiltration (Amicon® Ultra-15 Centrifugal Filter Units, MWCO 100 kDa). The concentrated HDL was purified based on the difference in their molecular size by size exclusion chromatography using a Superose 6 column (1.5×60cm) (GE Healthcare) Fast Protein Liquid chromatography [V_0_= 34ml, Flow rate= 0.4ml/min, fraction size= 1.5ml (more information on size exclusion chromatography can be obtained at https://www.gelifesciences.com/en/us/solutions/protein-research/knowledge-center/protein-purification-methods/size-exclusion-chromatography#size-exclusion). The TLF enriched HDL fractions that showed 50% or higher trypanolysis were collected and concentrated [11]. The concentrated fraction was again tested for trypanolysis to calculate lytic unit to confirm the enrichment of TLF in human HDL preparation. Therefore, we use human HDL interchangeably with TLF in the paper. The predominant lipoprotein fraction in bovine is high-density lipoproteins. Therefore, for the bovine HDL separation, following the first float after density gradient centrifugation (total lipoprotein), HDL was purified by size fractionation on a Superose 6 column (GE Healthcare) equilibrated with TBS. Only the fractions containing APOA1, the structural protein of HDL were then pooled and concentrated.

For mouse lipoprotein purification, mouse plasma was collected, and density was adjusted to 1.25 g/ml and lipoproteins (top fraction) was collected by density gradient centrifugation as described above. The lipoprotein was further purified from the KBr added to adjust density by dialysis followed by concentration as described above for human HDL. For separation of lipoproteins from plasma by size exclusion chromatography, 100□l of mouse plasma was separated based on their size using superdex 200 10/300 column, (GE Healthcare) (V_0_= 24ml, Flow rate= 0.5ml/min, fraction size= 1.0ml) in which larger molecules are eluted first followed by the ones that are smaller.

### Ammonium chloride neutralization and infection of macrophages

*L. major* metacyclic promastigotes were opsonized for 30 min in DMEM containing 4% A/J mouse serum. Bone marrow derived macrophages in DMEM culture medium, were infected at a multiplicity of infection of 3 parasites per macrophage for 2 hours at 33°C (5% CO_2_, 95% air humidity). Thereafter, non-phagocytosed parasites were washed off with prewarmed DMEM. Macrophages were then treated with 10 mM NH_4_Cl for 30 min to neutralize the acidic compartments of the cell before the addition of HDL (1.5 mg/ml TLF and bHDL). The cultures were further incubated in the presence or the absence of HDL for 24 hours. Intracellular parasites were quantified after staining with DAPI (3 µmol/L) by a Leica TCS SP2 AOBS confocal laser-scanning microscope.

### Changes in intracellular pH

Changes in intravacuolar pH were evaluated using Acridine Orange (AO) (A3568, Sigma). BALB/c bone-marrow derived macrophages were infected with *L. major* or *L. amazonensis* metacyclics for 2 hours as described above, before addition of TLF or bovine HDL (1.5 mg/ ml) or NH_4_Cl (10 mM). Infected and uninfected cells were then incubated with 10 µM AO in PBS for 10 min at room temperature. Samples were immediately analyzed using a Leica TCS SP2 AOBS confocal laser-scanning microscope.

### *Leishmania amazonensis in vitro* assay

TLF or bovine HDL (1.5 mg/ml, physiological concentration in blood) was added to parasites in Dulbecco’s Modification of Eagle’s Medium (DMEM, Corning Cellgro) with 0.2% Bovine Serum Albumin (BSA, Sigma) (Ficoll purified metacyclic promastigotes or axenic amastigotes) at neutral pH 7 (mimics extracellular environment) followed by acidification to pH 5.6 using succinate buffer (0.05M succinic acid and 0.015M NaOH, pH 3.8) to mimic phagosome. Parasites were incubated at 33°C (5% CO_2_, 95% air humidity) for 1 hour. Parasites with HDL were then subjected to second acidification to pH 4.5, (Increasing acidification, IA) or neutralization (Acid then Neutral, AN) to pH 7 and incubated for another hour at 33°C (5% CO_2_, 95% air humidity). To assess the number of remaining intact parasites, we used a hemocytometer.

### Parasite centrifugation and washes

Purified *L. amazonensis* metacyclic promastigotes or axenic amastigotes (1 x 10^6^/ml) were centrifuged (4^°^C, 8000 x g, 1 min) to obtain the pellet. For subsequent washes, parasites were then resuspended in DMEM, 0.2% BSA and centrifuged as described above.

### TLF/HDL labeling and binding to parasites

TLF and bHDL were labeled with DyLight™ 488 NHS ester (ThermoFisher, 46402) according to the manufacturer’s instructions. Purified *L. amazonensis* metacyclic promastigotes or axenic amastigotes (1 x 10^6^/ml) were washed twice at 4°C as described earlier and incubated with 10 μg/ml Dylight-488 labeled TLF/HDL in DMEM, 1% BSA for 30 min on ice. Cells were washed twice at 4°C as described above before being analyzed by flow cytometry. Flow cytometry was performed with a Becton Dickinson FACSCalibur system.

### Detection of surface glycoconjugates

To test for the presence of surface glycoconjugates, 2×10^7^ metacyclic promastigotes of wild-type, complemented lines (l*pg1*^*-/-*^+LPG1, l*pg5A*^*-*^*/lpg5B*^*-*^*/*+LPG5A/5B) and mutants (l*pg1*^*-/-*^ l*pg5A*^*-*^*/lpg5B*^*-*^) were lysed in 1% NP-40 in the presence of an protease inhibitor cocktail (Sigma, 11836170001). Equal cell equivalents were separated on a 10% SDS PAGE (Biorad, 4561036) gel followed by western blot using anti-phosphoglycan antibody WIC79.3 (1:1000; gift from Dr. Steve Beverley, WU), followed by anti-mouse (1:10,000, Proteintech, SA00001-1) secondary antibody and detected by chemiluminescence.

## Supplementary Figures

**S1. HGD transfection increases the myeloid cell count in the blood**

Mice were transfected by HGD and blood was collected 24 hours post-HGD (n=4) and stained with myeloid cell markers CD11b, Ly6G, and Ly6C and analyzed by flow cytometry. Untransfected mice were used as controls (n=2). The data represent myeloid cells from 1 mouse each.

**S2. Quantification of APOL1 and HPR protein in HGD and germline transgenic mice**

**A.** Baboon plasma was serially diluted (1:40 to 1:320) and ran on a non-reducing SDS PAGE gel along with mouse plasma collected by tail bleeding from targeted germline transgenic mice that was serially diluted (Homozygous mice-1:40-1:320 and Heterozygous mice-1:20-1:320). Separated proteins were then probed by western blot for baboon APOL1 and HPR. * Proteolyzed APOL1. **B.** Mouse plasma was collected by tail bleeding and diluted in SDS PAGE loading buffer (1:40 for transiently transgenic HGD mice injected with the “human plasmid” with *APOL1:HPR* and 1:10 for homozygous germline transgenic mice expressing baboon *APOL1* and *HPR*). The proteins were separated on a non-reducing SDS PAGE gel and probed by western blot for APOL1 and HPR. A known concentration of recombinant proteins (human APOL1 and haptoglobin for HGD and baboon APOL1 and human haptoglobin for germline transgenic mice) were used as standards to determine the concentration of the respective proteins. Quantitation of the protein band was performed using Image J software.

**S3. Neutrophil gating strategy and depletion from blood and ears in mice** Neutrophils were depleted from mice using 1 mg anti-mouse Ly6G clone 1A8 antibody (1A8) or an isotype IgG2A antibody (Isotype) **A.** Mouse blood (50μl) was collected by tail bleed 24 hours after antibody treatment (time of infection); **B.** Mouse ears were collected 10 hours after infection with 1×10^6^ metacyclic promastigotes and processed as described in the methods section. The white blood cells were then stained with anti-mouse CD11b PE, anti-mouse GR-1 FITC, and anti Ly6C APC and measured by Flow cytometry using a BD FACSCalibur™. Total cells were then sub-gated for CD11b^+^ lineage cells. CD11b^+^ lineage cells were then divided into sub-populations. Neutrophils were identified as the CD11b^+^Ly6G^+^GR1^+^ subpopulation.

**S4. TLF does not change the pH of macrophage phagosomes**

**A–D.** BALB/c bone-marrow derived macrophages were infected with *L. amazonensis* metacyclics for 2 hours before addition of 10 mM NH_4_Cl (**B**), 1.5 mg/ml TLF (**C**) or bovine HDL (**D**) for 24 hours. Infected macrophages were stained with Acridine Orange after 24 hours and fluorescence emissions of 530 and 640 nm were acquired simultaneously. **A.** untreated infected cells; **B.** NH_4_Cl treated cells; **C.** TLF-treated cells; and **D.** bovine HDL-treated cells. Images acquired using Leica confocal microscope. **A’-D’** phase contrast images of the macrophages in A-D.

**S5. Binding of *L. major* metacyclic promastigotes to TLF or bHDL**

*L. major* metacyclic promastigotes (1×10^6^/ml) were treated with 10 µg/ml of DyLight-488 labelled TLF and bHDL (blue) or not (red) for 30 min on ice. Fluorescence intensity was quantified by flow cytometry.

## Acknowledgements

We would like to thank Dr. David Sacks at National Institute of Allergy and Infectious Diseases, Dr. Gerald Spaeth at the Pasteur Institute, and Katherine L. Owens at Washington University School of Medicine for their technical advice for the *Leishmania* work; Dr. Arturo Zychlinsky at Max Planck Institute and Dr. Amariliz Riviera-Medina at Rutgers for their technical advice for neutrophil work. We would like to thank Julian Sherman for his technical assistance.

## Author Contributions

Conceived and experiments designed: JR,JP, MS, MTN. Experiments performed: JP, MS, MTN, MK, JV. Data Analyzed: JP, JR, SMB, MS, MTN. Wrote the paper: JP, MS, MTN, JV, MK, SMB, JR.

